# The costs and benefits of dispersal in small populations

**DOI:** 10.1101/2021.12.16.472951

**Authors:** Jitka Polechová

## Abstract

Dispersal has three major effects on adaptation. First, gene flow mixes alleles adapted to different environments, potentially hindering (swamping) adaptation. Second, it brings in other variants and inflates genetic variance: this aids adaptation to spatially (and temporally) varying environments but if selection is hard, it lowers the mean fitness of the population. Third, neighbourhood size, which determines how weak genetic drift is, increases with dispersal – when genetic drift is strong, increase of the neighbourhood size with dispersal aids adaptation. In this note I focus on the role of dispersal in environments which change gradually across space, and when local populations are quite small such that genetic drift has a significant effect. Using individual-based simulations, I show that in small populations, even leptokurtic dispersal benefits adaptation, by reducing the power of genetic drift. This has implications for management of fragmented or marginal populations: the beneficial effect of increased dispersal into small populations is stronger than swamping of adaption under a broad range of conditions, including a mixture of local and long-distance dispersal. However, when environmental gradient is steep, heavily fat-tailed dispersal will swamp continuous adaptation so that only patches of locally adapted subpopulations remain.

## Introduction

The role of gene flow in preventing (swamping) local adaptation is well acknowledged (Bürger, 2013; Yeaman, 2015). Dispersal between discrete niches will swamp adaptation when the respective fitness advantages in the distinct niches are not balanced, unless selection and/or niche-preference are strong (Barton, 2010; Bulmer, 1972; Haldane, 1930; Lenormand, 2002; Smith, 1970; Smith & Hoekstra, 1980). In his classic study, Haldane (1956) asserted that even in continuous space, gene flow could swamp adaptation, even if the nascent asymmetry in population density was small: the gene flow leads to departure from the optimum, and the resulting maladaptation translates to a lower density (assuming hard selection, where the fitness difference does not change with density or frequency of other types; Christiansen 1975): Through this positive (but detrimental) feedback, the asymmetry in population size would grow. The dynamics have been asserted to potentially form a stable species’ range margin, an idea also popularised by Mayr (1963). This idea appears supported by studies of adaptation to distinct niches with source-sink dynamics (i.e., highly asymmetric densities), which can be interpreted as adaptation to marginal habitats (Gomulkiewicz *et al*., 1999; Kawecki *et al*., 1997; Pisa *et al*., 2019). Yet, swamping of adaption is not generally expected in continuous space: even if the environment changes sharply, spatial clines in allele frequencies readily form and are further stabilised by coupling (evolution of linkage disequilibrium) with other clines, effectively strengthening the selection (Barton, 1979, 1983; Barton & Hewitt, 1985; Kruuk *et al*., 1999).

Whether gene flow swamps or benefits adaptation in continuous but spatially heterogeneous environments depends on further details. Classic theory predicts that when genetic variance is fixed (or constrained to evolve very slowly), dispersal across environments can indeed swamp adaptation (Kirkpatrick & Barton, 1997). As dispersal load (the incurred fitness cost of dispersal across environments on mean fitness) increases, continuous adaptation fails when genetic variance (measured by the variance load) is too small. Yet, when genetic variance increases with gene flow, the increase of variance aids adaptation such that it remains continuous, as long as the local density stays above zero (Barton, 2001). In small populations, the third effect of dispersal becomes important: the increase of the neighbourhood size (Wright, 1931). It has been shown that in two-dimensional habitats, gene flow can thus – perhaps counterintuitively – facilitate adaptation to environmental gradients (Polechová, 2018). This is because neighbourhood size rises with the dispersal distance squared, while the dispersal load only rises with the dispersal distance (as long as the environment varies mainly along one spatial dimension, such as along altitudinal or latitudinal gradients).

Yet, the assumption of Gaussian dispersal kernel (assumed in the theory above) may have been quite restrictive. It is conceivable that with a long-distance component to dispersal (as opposed to just a Gaussian dispersal kernel), the swamping effect would quickly overwhelm the beneficial effect of reducing genetic drift. While numerous studies stressed the importance of long-distance dispersal for maintenance of connectivity in fragmented populations (Davies *et al*., 2015; Kramer *et al*., 2008; Trakhtenbrot *et al*., 2005); theoretical exploration of the effect of long-distance dispersal on adaptation to environmental gradients has been limited. On the other end of the spectrum, with uniform dispersal across heterogeneous environments without a concept of local distance – such as in the island model – differential adaptation is readily prevented when the migration rate and the span of the environments is large enough, aided by asymmetry in selection and/or relative size of the niches (Szép *et al*., 2021).

## Model and Results

I focus on evolution of a species’ range in a two-dimensional habitat, assuming stabilising selection towards an environmental gradient *b*, which is stable through time. The trait *z* under selection is determined by a large number of additive loci (set to 500), and stabilising selection takes the form of *r*_*g*_(*z*) = −(*z* − *bx*)^2^*/*(2*V*_*s*_). Population growth is density-dependent *r*_*e*_(*N*) = *r*_*m*_(1 − *N/K*), where *K* gives the carrying capacity, and *N* the local population density. The mean fitness is 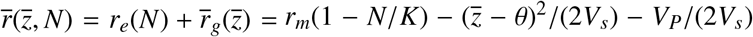. The term *V*_*P*_*/*(2*V*_*s*_) gives the load due to phenotypic variance *V*_*P*_ = *V*_*G*_ + *V*_*E*_, *V*_*s*_ is the variance of stabilising selection. In this model, one can use *V*_*G*_ ≡ *V*_*P*_ without a loss of generality: the loss of fitness due to environmental variance *V*_*E*_ can be included in 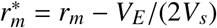, where 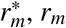 is the maximum per capita growth rate.

I assume the spatial optimum varies along one dimension (*X*); but it is constant along the other dimension (*Y*). The demes form a two-dimensional lattice of 100 demes in both dimensions. Mating is local within demes after migration and selection: the mating pool is given by (haploid) neighbourhood size 𝒩 = 2*πσ*^2^*N*. The trait is additive, and each bi-allelic haploid locus can contribute to trait value by the effect of *α* = 0.2. The population starts well adapted in the central half of the habitat: the optimum is matched by a series of clines spaced *α/b* apart, with cline width 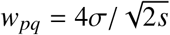, where *s* = *α*^2^*/*(2*V*_*s*_). In contrast to the model in (Polechová, 2018), dispersal is modelled as a mixture of two Gaussians, with two different variances *σ*_1_ and *σ*_2_, one with much higher than the other: (1 − *a*) *φ*_*N*_(0, *σ*_1_) + *a φ*_*N*_(0, *σ*_2_) Nichols & Hewitt (1994). As a default, *σ*_2_ = 20*σ*_1_. The sizeable asymmetry means that the proportion of long-range dispersal increases with *a* unless *a* is very close to 1, when all dispersal comes from the wider Gaussian.

While large populations will maintain clinal variation and gradually expand if the environment varies smoothly (Barton, 2001), adaptation across steep environmental gradients can fail abruptly when genetic drift is strong. As clinal variation dissipates, continuous adaptation fails and the species range collapses and/or fragments: each subpopulation is only adapted to a distinct optimum (Polechová, 2018). In contrast, continuous adaptation means clines form across the range, facilitating steady range expansion. Figure 1 shows that in small populations, adaptation across steep environmental gradients is rescued by increasing Gaussian dispersal (A). When the proportion of the long-range dispersal is small, adaptation is further facilitated, and range expands faster (B,C). However, as the proportion of long-range dispersal increases (D), gene flow swamps adaptation to steep environmental gradients, and continuous adaptation is no longer possible. This results in a population consisting of locally adapted subpopulations, with large gaps between them. This is demonstrated in Figure 2, which shows the spatial distribution of genetic variance as long-range dispersal increases. In the absence of long-range dispersal, genetic drift is too high for the spatial gradient, very few clines underlying adaptation are maintained, and local genetic variance is (mostly) low. As dispersal increases, genetic drift weakens and adaptation becomes easier: this is quite robust even when dispersal is leptokurtic (B,C). Eventually, though, increasing long-range dispersal swamps continuous adaptation, and the resulting population consists of locally adapted subpopulations spaced-apart by the long-range dispersal (D). Note that even broader dispersal can push the whole population to extinction as the mean fitness drops to zero globally. The parameters here are chosen so that this extreme is avoided; in general, it is dependent on the specific form of density-regulation.

**Figure 1:**
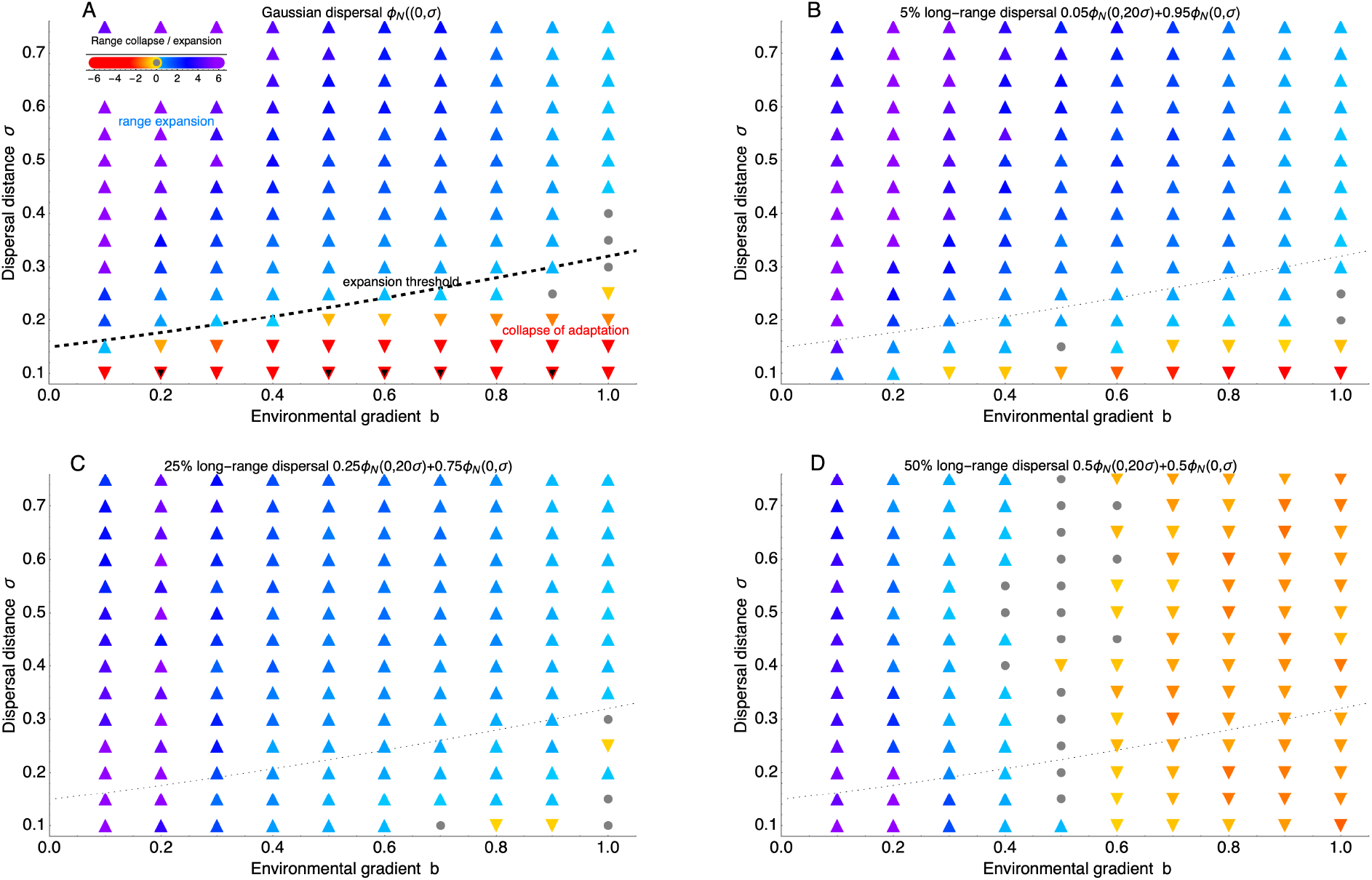
Increasing long-range dispersal first aids and then swamps adaptation across steep gradients. (A) In small populations, Gaussian dispersal aids adaptation to spatially varying optimum as the effect on reducing genetic drift is bigger than the cost of swamping: this allows the population to expand continuously along a gradient. Dispersal still aids continuous adaptation when a minor proportion disperses much further, so that the dispersal kernel is leptokurtic (B,C): with 5% resp. 25% of long-range dispersal. However, when the extent of long-range migration increases further (D, 50%), gene flow across steep gradients (*b >* 0.5) starts swamping adaptation. Dashed line (A) evaluates the expansion threshold under Gaussian dispersal 𝒩 ⪆ 6.3 *B* + 0.56, where the effective environmental gradient is 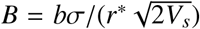 and the neighbourhood size 𝒩 = 4*πN*_*e*_*σ*^*2*^ *=* 2*πNσ*^*2*^ (Polechová, 2018). It is shown as a faint dotted line in pictures (B-D) for a reference; recalculation of the approximation of the threshold using the joint variance overestimates the rescue effect for longe-range dispersal (marginally out of the range for (B)). Parameters: local carrying capacity *K* = 4*r*_*m*_*/r\**, rate of return to equilibrium 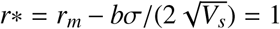, width of stabilising selection *V*_*s*_ = 1*/*2, mutation rate *µ* = 10^−6^, habitat width 100 demes along spatial gradient, 100 demes in the neutral direction, along which the optimum is fixed. The colour shows the rate of expansion (blue and purple hues) or indicates past or ongoing contraction (orange and red hues); black triangles denote populations which went extinct within 500 generations.

**Figure 2:**
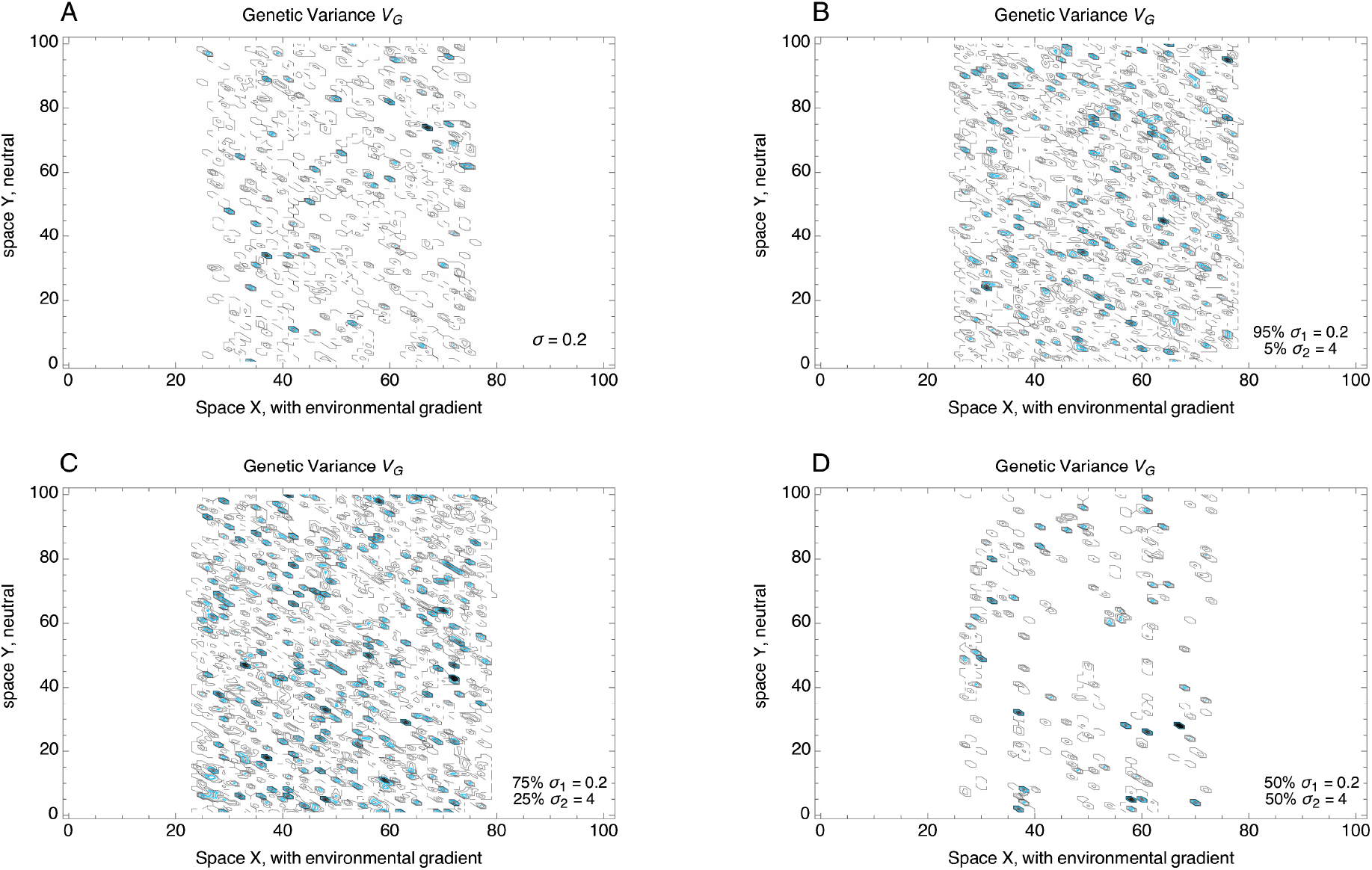
Illustration of spatial distribution of genetic variance for steep gradients, with increasing proportion of long-range dispersal - as in Fig. 1. (A) with only local dispersal, neighbourhood size is small and genetic drift overwhelms adaptation to steep environmental gradient: clines only form sparsely. The blue contour line depicts genetic variance of 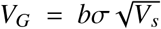, which would be maintained in the absence of genetic drift with Gaussian dispersal kernel *ϕ*_*N*_(0, *σ*) (Barton, 2001). Increasing long-range dispersal weakens the genetic drift, at first facilitating continuous adaptation across the species’ range (B,C). With strongly leptokurtic dispersal, however, continuous adaptation is swamped around singular, locally adapted populations (D). Because the environmental optimum does not change along the Ydimension, locally adapted sub-populations form stripes: gene flow is only realized along the neutral dimension. Parameters as in Fig. 1, *b* = 0.7; local dispersal ∼ *ϕ*_*N*_(0, *σ*_1_ = 0.2), long-range dispersal ∼ *ϕ*_*N*_(0, *σ*_2_ = 4).

## Discussion

The role of dispersal in spatially structured populations is complex: it can both swamp and facilitate adaptation (Garant *et al*., 2007). Additionally, in small populations, the effect of reducing genetic drift (and inbreeding depression) becomes important (Willi *et al*., 2006). Although often, the focus has been on how much of swamping by gene flow can still be balanced by selection (Bürger, 2013; Holt & Barfield, 2011; Yeaman, 2015), a number of studies have found that locally favourable alleles can be most readily established with intermediate migration supplying the new variants (Débarre *et al*., 2013; Gomulkiewicz *et al*., 1999; Uecker *et al*., 2014). This can be crucial for adaptation to both spatial and temporal variation. It has been suggested that for example in trees, variants brought in by long-distance pollen dispersal may aid adaptation to temporal change (Aguilée *et al*., 2016; Kremer *et al*., 2012).

The mixture of two dispersal kernels used in this paper can be seen as a model applicable to such a system, with two modes of dispersal: one more local (seeds), and one more global (pollen). Long-distance dispersal of gametes (such as planktonic transport of larvae), while adults disperse on a local scale, is also predominant in many marine organisms (Jordano, 2017; Kinlan *et al*., 2005). This note shows that in such systems, where dispersal is leptokurtic, gene flow across continuously heterogeneous environments does not easily swamp adaptation. Only extensive long-distance dispersal across steep environmental gradients does: population then fragments and only patches of locally adapted subpopulations remain. When population density is low, the benefit of gene flow in reducing the power of genetic drift, is more important. This can be relevant for management of marginal or fragmented populations: the oft-cited conclusion that gene flow swamps adaptation in marginal populations does not generally hold when genetic variance can evolve (Kirkpatrick & Barton, 1997). Interestingly, this is in line with a recent review which has found little evidence of gene flow swamping adaptation at the range margins (Kottler *et al*., 2021).

It is to be noted, though, that there is no quantitative meaning of a particular percentage of long-distance *vs*. local dispersal in this model. The chosen ratio of the widths of the dispersal kernels is largely arbitrary: if they were more similar, genetic drift would be reduced more effectively. There are several further limitations to the study: adaptation occurs via many loci of uniform and small effect, and hence it assumes weak selection per locus. It is possible that relaxation of this assumption would make pockets of adaptation more robust to the combined effect of genetic drift and swamping by gene flow. Secondly, it simulates small neighbourhoods, where genetic drift is strong and is thus much better suited to addressing adaptation at range margins than at central parts of a species’ range. While long-distance dispersal may still be beneficial in large populations (i.e., with a large neighbourhood size), for example by bringing in adaptive variants, such an effect is not addressed in this note.

